# Bridging biodiversity and gardening: Unravelling the interplay of socio-demographic factors, garden practices, and garden characteristics

**DOI:** 10.1101/2023.12.03.569783

**Authors:** Zsófia Varga-Szilay, Kinga Gabriella Fetykó, Gergely Szövényi, Gábor Pozsgai

## Abstract

The expansion of urban areas threatens biodiversity, disrupts essential ecological relationships and jeopardises fragile ecological networks, thereby impedes key ecosystem services. To avert irreversible consequences, there is a focus on improving the biodiversity value of domestic gardens for both human well-being and conservation and a global imperative for well-planned and sustainable urban environments. Here, we employ machine learning and network analysis and examine gardening practices and garden owners’ environmental consciousness in Hungary through a questionnaire-based study to untangle the interplay among socio-demographic factors, garden management, and garden characteristics. We found that the activities determined as biodiversity-positive were widespread among respondents, but a lack of undisturbed areas (n = 624, 49.52%), mowing several times a month (n = 404, 32.06%) and ubiquitous pesticide use (n = 783, 62.14%) were also present. Middle-aged respondents demonstrated more biodiversity-supporting activities than those over 55, who had long-term gardening experience and were predominantly conventional gardeners. Residents of towns showed the least biodiversity-positive activities, whereas those living in cities and the countryside fared better. Additionally, multiple interconnected garden characteristics revealed various types of gardens distinguished by care practices and use, such as gardens for food self-provisioning, ornamental gardens, or those prioritizing biodiversity support. Our results show that garden owners use pesticides, and within them herbicides, independently of socio-demographic parameters, gardening practices, or garden characteristics, suggesting a widespread pesticide use in Hungary.

Our findings suggest that strategies, to promote biodiversity-friendly gardening practices may not be equally suitable for all European countries with different cultural backgrounds, environmental consciousness and pesticide use. In particular, factors like differences between societal groups underscore the preference for in-person programs over online information transfer in several cases, for instance, among the elderly and those living in the countryside. This study offers fresh perspectives on the intricate connections between garden diversity, characteristics, and practices, and it lays the groundwork for future research into the sociological drivers of gardening practices in Eastern Europe. Our work also emphasises that optimizing gardens for multiple ecosystem services, including biodiversity conservation and enhancing well-being across diverse societal groups, requires a nuanced understanding of both ecological and socio-demographic factors.

**Highlights:**

- Biodiversity-friendly domestic gardens are key for urban sustainability
- Gardening practices of Hungarian gardeners varied with socio-demographic factors
- Lack of undisturbed areas, frequent mowing and pesticide use were the most harmful
- Elderly gardeners showed less biodiversity support and environmental awareness
- Tailored strategies are crucial for enhancing Eastern European garden biodiversity

## 1. Introduction

The expansion of urban areas poses an increasing threat to biodiversity, negatively affects crucial ecological relationships and threatens fragile complex ecological networks (Hagen et al., 2012), which, in turn, hamper ecosystem services such as pollination and biological pest control, and ultimately human well-being (Jabbar et al., 2022). In order to avoid reaching a state when the result of these effects are irreversible, there is a global need for better better-planned and sustainable urban environment (Breuste et al., 2013; Heidt & Neef, 2008; Ramalho & Hobbs, 2012). One important aspect of these needs is improving the quality and quantity of urban green spaces, both for human well-being and biodiversity conservation (Baldock, 2020). Indeed, in some developed countries, the number of urban green areas (such as public parks, green roofs, and community and private gardens) is increasing (Kabisch & Haase, 2013) and their potential in multi-purpose sustainability development is increasingly recognised. Although 16 to 27% of urban green spaces in Europe belong to private owners (Goddard et al., 2010), the importance of private or shared gardens (e.g. community gardens) in influencing the quality of urban green ecosystem is often underestimated (Camps-Calvet et al., 2016).

Domestic/home gardens and allotments are green spaces in urban ecosystems where people usually cultivate various plants (mostly fruits, vegetables and ornamental plants), and areas they use for recreation, outdoor activities, or even to connect with nature (Bell et al., 2016). Indeed, as a popular pastime, gardening is beneficial to mental health and strengthens well-being (Krols et al., 2022). On the other hand, domestic gardens also have a great potential as habitat refuges for wildlife and they can strongly contribute to maintaining high biodiversity (Cameron et al., 2012).

The first studies aiming to detect the biodiversity of gardens in Western Europe and North America were conducted at the beginning of the 1990s (Delahay et al., 2023). Interest in the role of urban and suburban gardens in the preservation and support of urban ecosystems has increasingly gained interest since then (Delahay et al., 2023) and showed that even small, but resource-rich garden habitats can significantly increase insect diversity (Griffiths-Lee et al., 2022) and support ecological services such as pollination, biological pest control or climate regulation (Andersson et al., 2007; Cavan et al., 2021). Gardens can function as ecological corridors or stepping stones for a multitude of organisms in the otherwise barren and often hostile urban landscape and, particularly when they have favourable habitat features (e.g. ponds (Hill et al., 2021)), can become local biodiversity hotspots (Baldock et al., 2019; Prendergast et al., 2022).

However, the true conservation potential of domestic gardens is governed by garden care practices; intensive garden management can negatively impact garden diversity as well as the gardens’ environment (Fontaine et al., 2016; Lerman et al., 2018). There is a proven link between environmental degradation, the decline of the abundance and diversity of birds and insects, and environmentally aggressive gardening practices (Muratet & Fontaine, 2015; Tassin de Montaigu & Goulson, 2023a), such as the uncontrolled use and overuse of pesticides. The use of neonicotinoid-based insecticides, and herbicides containing glyphosate is of particular concern for insect biodiversity in gardens (Tassin de Montaigu & Goulson, 2023a, 2023b). Additionally, through spillover effects, the excessive chemical fertiliser use (Law et al., 2004), the frequent irrigation (Egerer et al., 2018; Fernández-Cañero et al., 2011), as well as the introduction of potentially invasive ornamental plants (Süle et al., 2023) contribute to environmental degradation of not only the garden but also the adjacent areas.

Gardens can also provide ecosystem disservices, for instance, the increase of pests or disease-carrying insects (such as mosquitos and ticks) (Yang et al., 2019), or wildlife causing fear or aversion (Baker et al., 2020; Soulsbury & White, 2015). Nonetheless, increased ecological knowledge stimulates environmental stewardship, and garden owners’ access to and effective trialling of biodiversity-friendly practices can alter public perceptions surrounding contentious issues, such as pesticide phase-down or biodiversity-friendly plant protection (Barthel et al., 2010). Moreover, recent studies highlighted a central role of garden owners’ attitudes and consciousness in either promoting or impeding wildlife-friendly gardening. Wildlife-friendly gardening is a multifaceted issue which is influenced by several factors, such as demographics, socio-demographic drivers, motivations for gardening (García-Antúnez et al., 2023; Philpott et al., 2020), and appreciation for nature (Clayton, 2007), as well as trust in environmental associations and access to biodiversity-related information (Coisnon et al., 2019) and various management practices (Goddard et al., 2013).

For this reason, it is key to gain a mechanical understanding of how gardeners’ cultural background, their aim for gardening, and their connection with the natural environment in their garden can affect their willingness for optimising gardening habits and garden management for conservation benefits.

However, biodiversity-friendly gardening and gardening practices are intertwined with the type and layout of the garden (Hanson et al., 2021), as well as with the presence or absence of certain plants (Paker et al., 2014, Salisbury et al., 2015) and this interconnected system is likely to be further shaped by socioeconomic factors (Hoyle et al., 2017). Little research has been done so far to untangle the joint impact of linked gardening practices, garden characteristics, socio-demographic parameters, and motivations for gardening, and it also remains unclear how tightly garden and garden owner characteristics are linked, and which combination of these, leads to biodiversity-friendly gardening practices.

Yet, if these were to show clear patterns, they could provide an easy means to assess the gardens’ potential for supporting biodiversity or whether they could serve as parts of a habitat network that supports biodiversity. With this information, specific recommendations could also be suggested to guide favourable modifications in gardening habits, leading garden owners toward environmentally friendly practices.

Thus, in this study, we use a combination of methods of machine learning and network analysis to investigate how gardening practices, motivation for gardening, and garden characteristics can influence biodiversity-friendly gardening. We paid particular attention on the interlinked characteristics of gardens and gardening practices and examined how socio-demographic factors influence biodiversity-positive or -negative practices. Nonetheless, our ultimate goal is to explore pathways for maximizing conservation benefits and, at the same time, maintaining or improving human well-being linked to domestic gardens and gardening.

We focused our work on Hungary, a country characterized by conventional gardening practices in home gardens, widespread in Eastern European countries, including excessive use of pesticides (Varga-Szilay & Pozsgai, 2022).

## 2. Material and methods

### 2.1. Questionnaire Design

We distributed an online questionnaire that consisted of 58 questions, with all but 4 questions requiring mandatory responses. These questions were organized into nine sections, covering: (i) garden location, (ii) socio-demographic parameters, (iii) garden characteristics, (iv-v) motivation and gardening practices, (vi) garden cultivation, (vii) pesticide usage, (viii) observed insects (mostly pollinators) in the garden, and (ix) closing questions. The questionnaire was designed in Google Forms and it took 10 to 12 minutes to complete.

All responses were recorded anonymously, however, respondents could provide their email addresses. Respondents could indicate their education on a four-, their gender on a three-level scale (male, female, other), and their residency at a county-level (NUTS3, https://ec.europa.eu/eurostat/web/nuts/background, accessed 30th November 2023).

The questionnaire was actively distributed between the 26th of October 2022 and the 1st June 2023 in Hungary. The questionnaire was distributed through gardening-related websites, various social media platforms (including Facebook and Instagram), and mailing lists. Additionally, we reached out for professional bodies, non-governmental organizations, gardening- and biodiversity-protection-related foundations, societies and organizations via email. Moreover, we used QR codes and hashtags to extend sharing efficiency.

As a general term, ‘pesticides’ were defined as all synthetic and non-synthetic products that are used to control pests. We included in the term all commercially available and homemade plant protection products, either those allowed in organic gardening/farming or used in conventional practices. The terms ‘pesticide’ and ‘plant protection products’ were used as synonyms. However, for the analysis, of the three recorded pesticide categories that of synthetic products was used separately. Although herbicides are also pesticides, because of the importance of some (e.g. glyphosate as a widely used herbicide in the agriculture and gardening practices (Benbrook, 2016)) we separately recorded whether garden owners used them (please see the Supplementary Table 1). Furthermore, the terms ‘domestic garden’, ‘house garden’, and ‘home garden’ were used as synonyms and the term ‘gardening’ included all garden work and all garden care practices, such as the cultivation of flowers, fruits, vegetables, and ornamental plants, mowing, and soil management.

In our interpretation, unmown patches refer to areas which otherwise would be mown but garden owners (intentionally) avoid mowing them, while undisturbed areas/fallow were permanently undisturbed areas that are independent of mowing. Although the responses about whether garden owners know plant, insect or bird diversity in their garden were self-evaluated, and thus subjective, we keep referring to them using the absolute term ‘knowledge’.

The No Mow May campaign (NMMc) urged garden owners not to mow their lawns in May, as this is the month with the most abundant food sources for pollinators (Plantlife’s No Mow May Movement, https://www.plantlife.org.uk/campaigns/nomowmay/, accessed 30th November 2023) in the Northern Hemisphere. Since this campaign was adopted in Hungary, translated as ‘Vágatlan Május’, and widely promoted across the country, as well as its biodiversity-friendly approach had a clear relevance to our study, we asked the respondents about their knowledge/participation.

The question assessing garden pollinator diversity was picture-based, with images illustrating representative members of pollinator groups commonly found in gardens.

### 2.2. Data processing

For the analysis, we used 27 questions relevant to our aims of the original 58 ones. The original categorical responses were on a few occasions re-categorised for analytical purposes (for details, please see Supplementary Table 1). For instance, the answers of ‘My favourite hobby’, ‘A pleasant pastime’, and ‘Opportunity to exercise’ were merged into ‘Pastime’ and the ‘Duty’ and ‘Work’ original levels into ‘Duty’. Respondents under the age of 18 (n = 5) were excluded from the analysis.

Due to methodological constraints, for parts of the analysis we used binarized data (see below). Some of our questions inheritably had binary responses (Yes/No) but those with multiple choices were converted to binary variables by providing a separate data variable for each choice. Since they provide little information, yet burden computational processes, variables in which response agreement was over 95% were removed for the co-occurrence analysis (see below).

### 2.3. Statistical analysis

To investigate how garden characteristics and gardening practices were influenced by socio-demographic factors, a distance-based redundancy analysis (db-RDA) was conducted by using the binarized responses as dependent variables and gender, age, education level, whether the garden owners had children, and whether they lived in a city, town, or countryside, as explanatory variables. Distances were calculated using the Jaccard distance measure. The significance of the model, the axes, and the variables were tested using an ANOVA-like permutational test for Constrained Correspondence Analysis (CCA), with 999 iterations.

In order to assess the major garden characteristics and socioeconomic factors driving biodiversity-positive and biodiversity-negative gardening practices, we selected six and five (respectively) proxy variables to represent the extremes of these habits. The positives included the active support of pollinators either by sources of water or nectar- and pollen-rich flowers (1); the active support of pollinators by natural (for example wildflower strips) (2); or artificial habitats (insect- or bee hotels, hoverflies-lagoons) (3); leaving unmown patches (4); having a pond (5) and complete avoidance of pesticides (6). The negative ones included the use of synthetic pesticides (1), (additionally) herbicides (2) or synthetic fertilisers (3), having no undisturbed patches (4), and mowing the lawn several times a month (very often) (5). For each respondent, both ‘good’ and ‘bad’ attributes were counted, each of these values were rescaled to between zero and one and the biodiversity-negative values were set to their corresponding negative values. The sum of these values (biodiversity friendliness score, BDF score, henceforth) was used as the response variable for Gradient Boosting Machine (GBM) learning processes.

Highly subjective questions, such as garden owners’ perceptions of their gardens as pollinator-friendliness and their willingness to participate in a garden network that helps maintain biodiversity, were excluded. All other garden characteristics, gardening practices, and socioeconomic factors, but those from which we calculated the BDF score, were included in the model as explanatory variables. For instance, since synthetic pesticide use was included among the biodiversity-negative practices, the variable coding pesticide use was excluded (see Supplementary Table 6 for the full list). After an optimalisation process (see code on https://github.com/zsvargaszilay/gardening_in_hungary), we fit the GBM model using a Gaussian distribution in 3 levels of interaction depth, with 0.1 shrinkage and 0.80 bag fraction on 85 trees. The model fit was evaluated by calculating the R-squared and root mean standard error (RMSE) values.

To improve interpretability, we used the SHapley Additive exPlanations (SHAP) of our GBM model. SHAP comprehensively assesses individual variable contributions, by considering variable interactions, assigns importance values and ensures fair comparisons through evaluations in all possible variable orders (Lundberg & Lee, 2017). For modelling and the visualization of model results, we used the ‘*gbm’* (Greenwell et al., 2022, version 2.1.8.1), ‘*caret’* (Kuhn et al., 2023, version 6.0-94), ‘*shapviz’* (Mayer & Stando, 2023, version 0.9.2) and ‘*kernelshap’* (Mayer et al., 2023, version 0.4.0) R packages in an R environment (R Core Team, 2021, version 4.3).

A probabilistic co-occurrence model analysis was used to investigate which responses were associated to each other. The analysis identifies pairwise associations based on the comparison of observed frequencies against those anticipated by chance. A positive association is inferred when the observed frequency exceeds the expected baseline and, conversely, a negative association is concluded when observed frequencies fall below the expected values (Veech, 2013). We considered positive associations as synergies and negative ones as antagonistic effects (trade-offs) in shaping gardening practices. Only associations with probabilities over 0.6 were considered in the final networks. Thus, the network consisted of garden characteristics and gardening practices as nodes and the associations between them as links. We examined the interdependencies in the association network and recorded the answers which were the most connected to other answers (i.e. had the most links to other nodes in the network). Groups of densely associated responses (modules) were detected by the Louvain community detection algorithm (Blondel et al., 2008).

In order to identify the major socioeconomic factors driving these associations, we conducted a redundancy analysis (RDA) using the similarity of subnetworks built for varying socio-demographic backgrounds. First, we generated all unique combinations of the responses of five socio-demographic variables: 1) the respondent’s gender (female, male, other); 2) age in three categories (younger than 36 years, between 36 and 55, and over 55); 3) whether the respondent lived in a city, a town, or in the countryside; 4) the respondent’s highest level of education (middle and high), and 5) whether they have children. This yielded 108 unique combinations (e.g. a younger than 36-year-old city-living female, with middle-level of education and children), which we individually used to query our dataset. If the query yielded at least 30 respondents, we used the same method as above to build association sub-networks. A matrix of either the positive (+1) or negative (-1) sign or the absence (0) of the pairwise associations in each query was used to conduct the RDA, with the five socio-demographic factors as explanatory variables and Bray-Curtis distance measures. An ANOVA-like permutation test for CCA, with 999 permutations, was conducted to test the model significance, the significance of each canonical axis and that of the explanatory variables.

Preliminary data clean-up was done in a Python (Python Software Foundation, 2019, version 3.8) environment, with the help of ‘*NumPy*’ (Harris et al., 2020) and ‘*Pandas*’ (The pandas development team, 2022) libraries. All further data manipulation, analysis, and visualisations were conducted in R version 4.3, with the help of the ‘*cooccur’* (Griffith et al., 2016), ‘*dplyr’* (Wickham et al., 2023, version 1.1.2) ‘*ggplot2’*(Wickham, 2016), ‘*igraph*’ (Csardi & Nepusz, 2006, version 1.5.1), and ‘*vegan*’ (Oksanen et al., 2022, version 2.6-4) packages.

## 3. Results

### 3.1. Socio-demographic drivers

Of the 1260 people who completed the questionnaire, 343 were males (27.22%), 916 were females (72.70%), and one person’s gender was not binary (0.08%). The majority of respondents were between 36 and 55 years old (53.10%). More garden owners with high education completed the questionnaire (n = 906, 71.90%) than those with middle-level education (n = 385, 28.10%), and almost half of the respondents had children (n = 616, 48.89%). The questionnaire was mostly filled out for domestic gardens larger than 500 m^2^ (n = 580, 46.03%) (for details of socio-demographic characteristics of the study population, please see the Supplementary Table 2). The willingness to respond was slightly unbalanced, with more respondents from the Western than from Eastern counties (Supplementary Figure 1). In terms of population, Zala county (HU223) was the most responding region (2.8 respondents for 100,000 inhabitants), and Szabolcs-Szatmár-Bereg county (HU323) was the least.

The constrained variables of the db-RDA model explained 3.72% of the total variance. The first and the second db-RDA axes explained 45.25% and 23.65% of the filtered variance, respectively (Figure 1). The model (p ≤ 0.001) and all explanatory variables were significant (for details see the Supplementary Table 3). Although most garden characteristics and gardening practices were located close to the origin (indicating a low correlation), some were highly correlated with one or two explanatory variables. For instance, living in the countryside explained the large garden size and house garden type, and the age over 55 correlated positively with gardening for a long time. Furthermore, middle age and having children influenced similar garden characteristics (e.g. growing vegetables and having undisturbed areas) and gardening practices (e.g. producing crops for consumption). The same variables were negatively affected by living in town and having a middle-level education.

**Figure 1:**
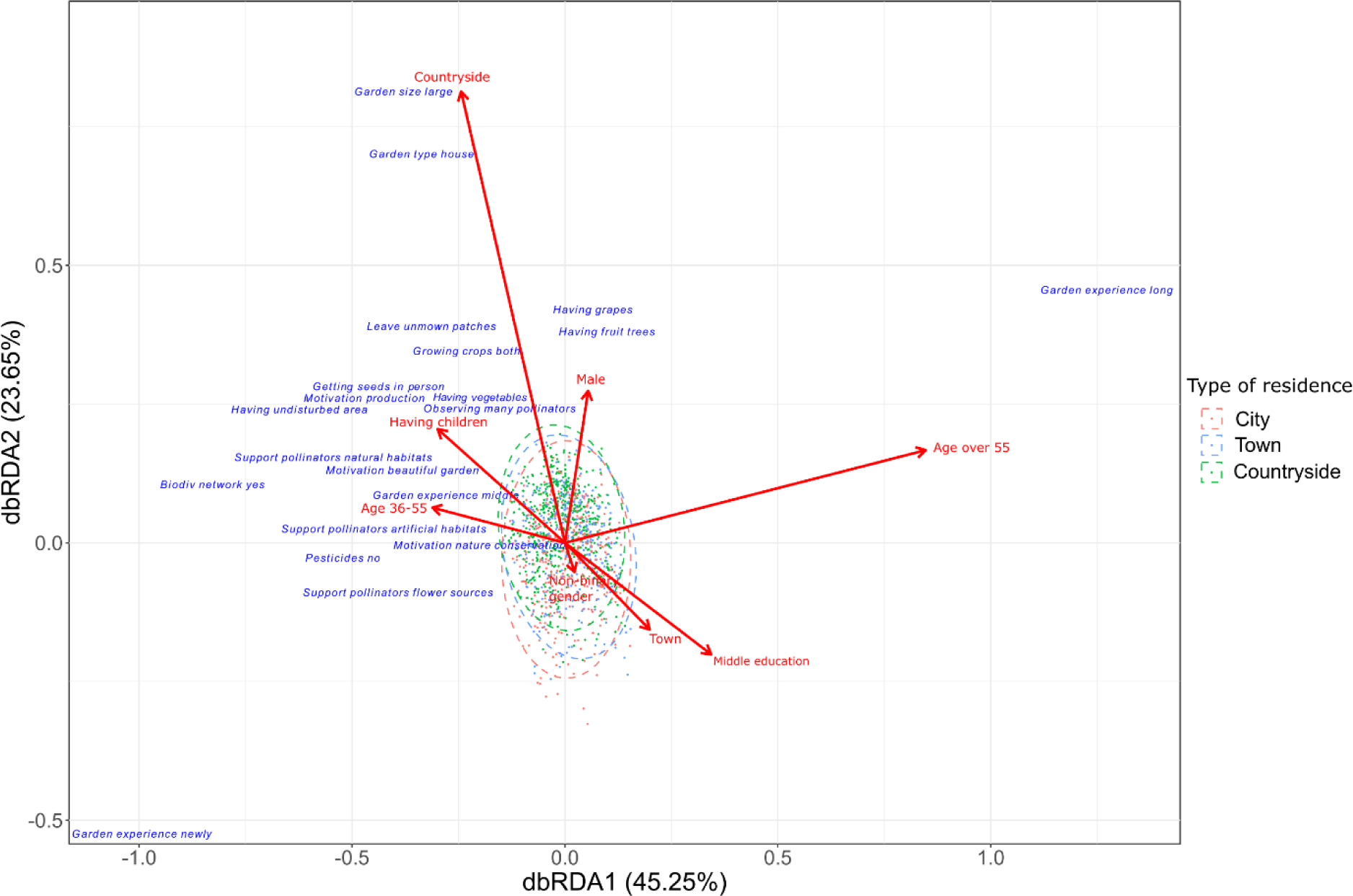
Distance-based redundancy analysis (db-RDA) plot showing the relationship among garden characteristics, gardening practices and socio-demographic (explanatory) variables. The length and direction of the vectors represent the strength and direction of the relationship. The ellipses represent the 95% confidence interval of the groups with different types of residency. Only garden characteristics and gardening practices (blue) whose RDA scores on the first two axes were lower than -0.2 or greater than 0.2 are shown.

### 3.2. Biodiversity-positive and -negative gardening practices

In the study population, the most common activity that had a biodiversity-positive effect in gardens was the supporting pollinators with water sources (n = 870, 69.05%) (Figure 2). This was followed by the supporting pollinators with pollen- and nectar-rich plants (n = 828, 65.71%) and establishing natural habitats (n = 827, 65.63%). Although a relatively large proportion of the respondents leave unmown patches when mowing (n = 786, 62.38%), only 37.86% (n = 477) of the respondents avoid pesticides completely. Neither creating artificial habitats (such as bee hotels) (n = 423, 33.60%) nor joining the NMMc (n = 284, 22.54%) were common activities among the respondents. Only 184 respondents (14.60%) had a pond (Supplementary Table 4A).

**Figure 2:**
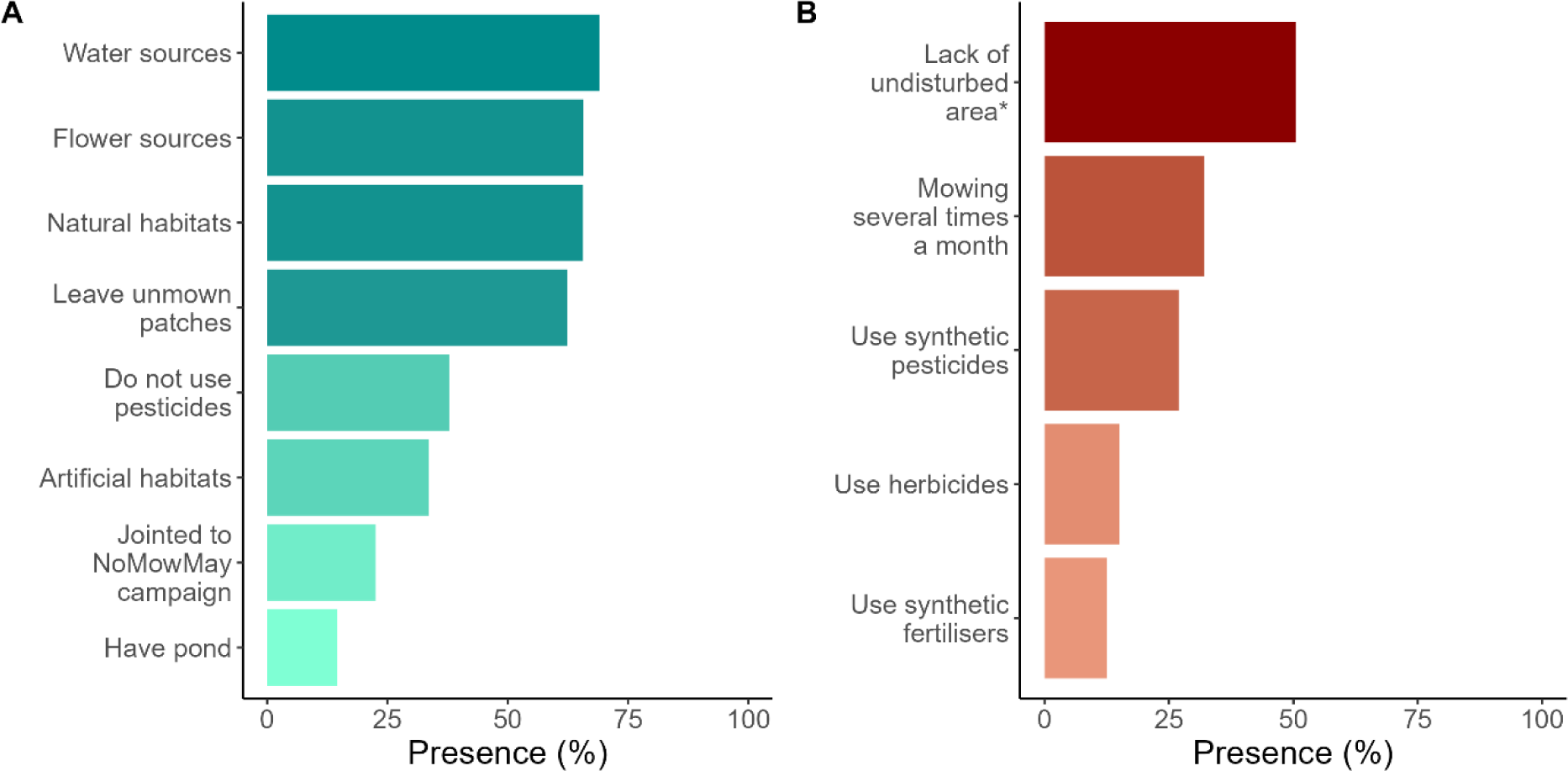
Presence (%) of the biodiversity-positive (A) and biodiversity-negative (B) gardening practices among respondents (n = 1260). Since to the question of whether respondents had undisturbed areas (*) the ‘no’ answer was considered as a biodiversity-negative practice, in this figure, the percent of ‘no’ answers is shown.

Among the gardening activities that may negatively impact domestic gardens’ biodiversity, the lack of undisturbed areas (624 respondents, 49.52%) was the most common, followed by mowing several times a month (n = 404, 32.06%). Synthetic pesticides were used by 26.98% (n= 340) of the respondents, whilst herbicides (n= 190, 15.10%) and synthetic fertilisers (n = 158, 12.54%) were less often used (Figure 2, Supplementary Table 4B). Of the herbicide users, 68.94% use glyphosate-containing herbicides (Supplementary Table 5).

The GBM model suggested that the best indicators for predicting whether gardening practices can be considered as biodiversity-positive or biodiversity-negative were the number of pollinator groups garden owners usually observe (relative influence: 11.68) and the type of the gardens, such as village garden or kitchen garden (relative influence: 9.14) (Supplementary Table 6). The model explained 16.00% of the variance of the dataset with the RMSE value of 0.32. The SHAP analysis determined that the information through personal links was the most important variable (SHAP value = 0.038), followed by the garden type (SHAP value = 0.032) (Figure 3). Variables, such as information through personal links, kitchen garden and orchard, knowledge about insects, less than 10 years duration of gardening experience, and middle age (36-55) shifted the outcome toward the biodiversity-positive direction. Having a flower garden and house garden, observing few pollinator groups in the garden, medium garden size (100-500 m^2^), more than 10 years of duration of gardening experience, living in town, and lack of herbs in the garden drove the outcome toward the biodiversity-negative direction.

**Figure 3:**
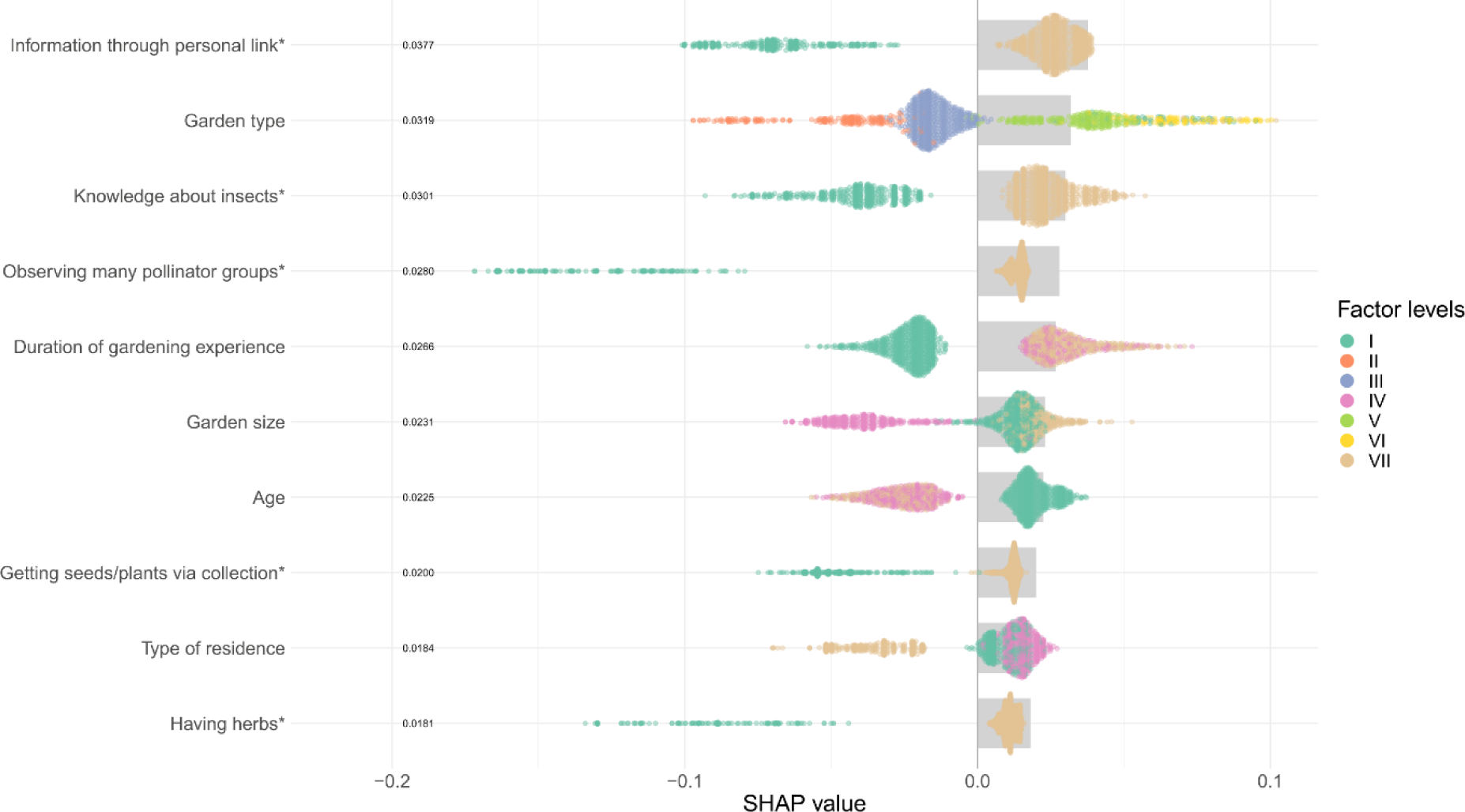
Global SHAP (SHapley Additive exPlanations) summary plot for the GBM model. The y-axis shows study variables ordered by their importance at a global model level by studying average absolute SHAP values as a bar plot (marked with grey colour). The y-axis also shows the variables’ importance by beeswarm plots of the SHAP values where each points represents one respondent, and colour represents the variable levels (Roman numerals in brackets indicate the factor levels). Variables marked with asterisks (*) have two levels (I and VII). The colour code for non-binary variables is as follows (from up to down): Garden type: Community garden and other (I), Flower garden (II), House garden (III), Kitchen garden (V), Orchard (VI), Vineyard (VII); Duration of gardening experience: Long ago (I), Middle (IV), Newly (VII); Garden size: Large (I), Medium (IV), Small (VII); Age: 36-55 (I), Over 55 (IV), Under 36 (VII); Type of residence: City (I), Countryside (IV), Town (VII).

### 3.3. Associations among garden characteristics, gardening care practices, and motivations of garden owners

Whether or not herbs were present in the garden had the most associations (i.e. links) in the network (15 positive and 1 negative) and the one garden owners considering their garden as pollinator-friendly was the second most connected with both positive and negative associations (14 and 1, respectively). The use of non-synthetic fertilisers had the same number of positive associations as the most connected node. Among the examined parameters, the lack of awareness of the NMMc exhibited negative associations with five variables (Figure 4).

**Figure 4:**
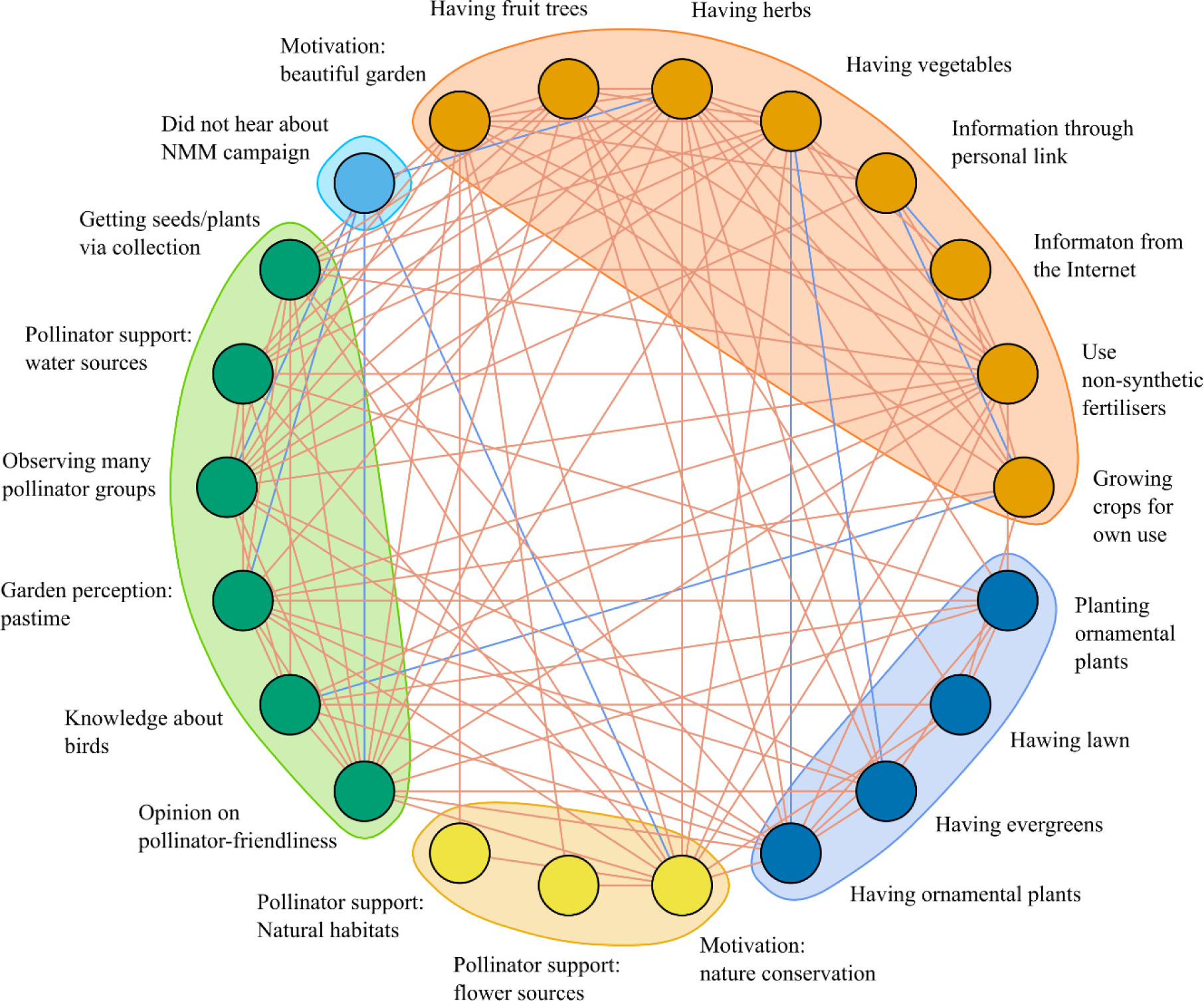
Co-occurrence network illustrating the statistically non-random associations among garden characteristics, gardening care practices, and motivations of garden owners. Red lines indicate positive, and blue lines negative associations. Variable colouring and shaded areas indicate modules identified by the Louvain community detection algorithm.

The community-detection algorithm identified four large modules and a module solely containing the variable of the lack of awareness about the NMMc. The largest group comprised of having fruits, vegetables, and herbs in the garden, along with a positive connection to a desire to make gardens beautiful, cultivating for own use and the use of non-synthetic fertilisers (Figure 4). Within this group, the means of gathering information through personal channels and from the Internet were negatively associated. Outside of the module, this group was negatively associated with the lack of awareness about the NMMc, having ornamental plants and evergreens, and the knowledge of birds. The second largest module contained variables such as observing high pollinator diversity, perceiving gardening as a pastime, and the knowledge of birds. This module was negatively associated, outside of the module, with the cultivation for own use and the lack of awareness about the NMMc. The third largest module, containing variables such as the planting of ornamental plants and the presence of lawns, evergreens, and ornamental plants in the garden, showed numerous positive associations both within and outside of the group, and one negative with having vegetables in the garden, from the largest group. The fourth module contained only three variables: two types of pollinator support (providing natural habitats and flower sources) and enthusiasm about nature conservation. These showed positive associations with each other and were negatively associated with the lack of awareness about the NMMc.

Of the variables we investigated, some, such as those related to pesticide or herbicide use, were not associated with any other variables (and thus did not appear in the network), indicating that the frequency of their co-occurrence did not deviate from that predicted by random chance.

The constrained variables of the RDA model explained 3.35% of the total variance with the first and second RDA axes explaining 28.27% and 19.91% of the filtered variance, respectively (Supplementary Table 7). Only the first axis was significant (p = 0.037). The majority of the connections did not separate well and were grouped near the origin. The associations between using non-synthetic fertilisers, having herbs in the garden and motivation for making the garden more beautiful, as well as that between collecting seeds and/or plants and observing many pollinator groups in the garden, correlated with the socio-demographic variable of the age over 55. Living in towns and having both ornamental plants and evergreens were also correlated. Of the explanatory variables, education was significant (p = 0.035), and the type of residence was marginally significant (p = 0.060). Indeed, there was little overlap between middle and high education levels in the RDA, whereas the confidence ellipse of respondents living in cities almost completely overlapped with those living in town or the countryside (Figure 5). Respondents under 35 were not separated from the other two age groups. However, the middle-aged (35-55 ages) and the respondents over 55 were separated from each other (Supplementary Figure 2).

**Figure 5:**
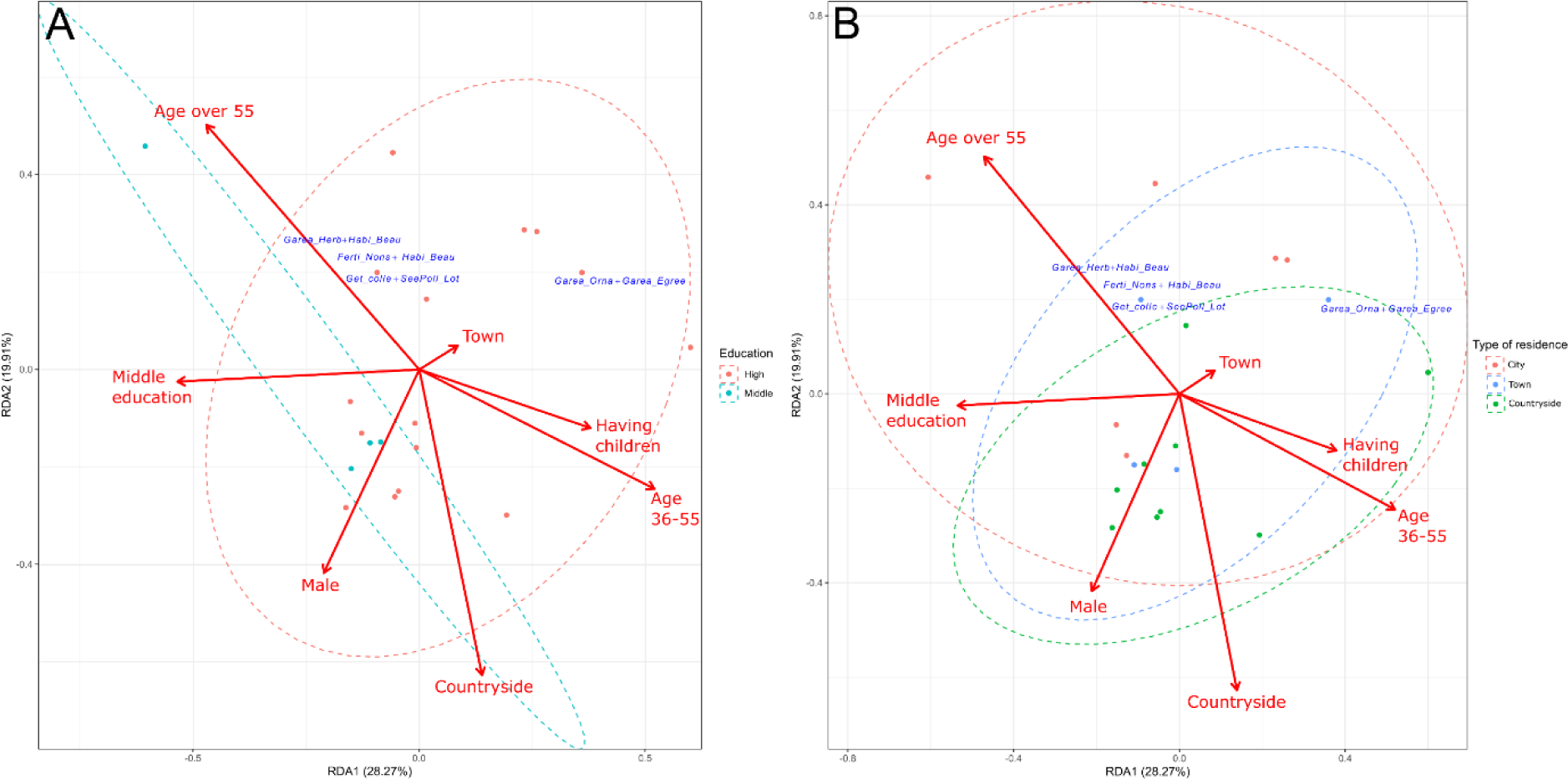
Redundancy analysis (RDA) plot showing the associated pairs of gardening practices and garden characteristics, along with the explanatory socio-demographic variables. The length and direction of the vectors represent the strength and direction of the relationship. The ellipses represent the 95% confidence intervals of associations of education levels (A) and resident types (B). Only associations (blue) whose RDA scores on the first two axes were lower than the mean of zero and the smallest value on the axes or greater than the mean of zero and the greatest value on the axes are shown. (Abbreviations: Garea_Herb: having herbs, Habi_Beau: motivation: beautiful garden, Ferti_Nons: use non-synthetic fertilisers, Get_colle: getting seeds/plants via collection, SeePoll_Lot: observing many pollinators groups, Garea_Orna: having ornamental plants, Garea_Egreev: having evergreens.)

## 4. Discussion

In this study, we distributed an online questionnaire in Hungary to investigate how gardening practices, motivation for gardening, and garden characteristics can influence biodiversity-friendly gardening, particularly focused on the interlinked characteristics of gardens and garden care practices and examined how socio-demographic factors influence biodiversity-positive and -negative practices.

We found that various socio-demographic parameters influence gardening practices and several garden characteristics were interlinked with means of garden care. Our results suggest that middle-aged respondents with higher education levels tend to choose biodiversity-positive activities, potentially due to their greater awareness and sensitivity to environmental issues (De Silva & Pownall, 2014; Meyer, 2015). Moreover, the high proportion of this group having children may also indicate that these are conscious decisions. This group can easily embrace novel, biodiversity-friendly management practices and therefore involving them into sophisticated conservation gardening and online citizen science projects is likely to be highly rewarding.

On the contrary, respondents over 55 demonstrated less or no support for pollinators, used more pesticides, and had less interest in nature conservation. This age group had a long (more than 10 years) gardening experience, probably with conventional garden care practices in Hungary. Indeed the use of non-synthetic fertilisers (manure is widely available in Hungary) and the collection of seeds and plants are all representative to this group. The importance of making their garden ‘more beautiful’ may however be misleading since traditionally beautiful gardens are often extensively managed and lack natural features (Burr et al., 2018). Of this age group 40% live in the countryside, where biodiversity is relatively high and conservation efforts can be particularly fruitful. Therefore engaging these garden owners in biodiversity-friendly practices would be of great conservation merit. This becomes especially crucial given the ageing demographic of the Hungarian society (Obádovics & Tóth, 2023).

Although, living in towns scored the worst for biodiversity-positive gardening, residing in cities does not negatively impact examined gardening activities, suggesting a diverse array of gardens and garden owners within highly urbanised areas. The similarities between the associations among gardening practices and garden characteristics in cities and the countryside imply the presence of traditional house gardens and less flat-based housing than in most Western European cities (European Commission, Eurostat, https://ec.europa.eu/eurostat/cache/digpub/housing/bloc-1a.html, accessed 30th November 2023). This lack of a stark urban(city)-rural(countryside) divide may stem from the historical development of several Hungarian cities evolving from agriculturally oriented rural areas (Beluszky & Győri, 2005) resulting in gardens with mixed uses, especially in the suburban zones. Recent trends also indicate a rise in mixed-use gardens driven by financial considerations, with individuals seeking self-sufficiency for higher quality and affordability. The distinctions in gardening motivation (Jehlička et al., 2020; Smith & Jehlička, 2013), the frequency of food self-provisioning (Alber & Kohler, 2008; Vávra et al., 2018) and pesticide use (Coisnon et al., 2019) between Western and Eastern Europe underscore the necessity for distinct approaches when aiming to enhance urban biodiversity in populations residing in cities and emphasise the need for studies from Eastern European countries similar to those already available for the West.

However, our results show that gardening practices and garden characteristics are equally influenced by all socio-demographic parameters. Thus, identifying one particular group of garden owners whose habits are highly biodiversity-negative, and therefore whose gardening practices should be altered to improve biodiversity, proves challenging solely on socio-demographic parameters. Unless a study utilizing a broader set of social-demographic factors manages to pinpoint this specific target group, conservation actions must be directed towards a broader segment of society.

Our results uncovered numerous (positive and negative) associations between garden characteristics and garden care practices. The grouping, based on the non-random, positive associations of the variables, suggests four well-separated approaches in garden care practices and garden use. Gardens with a predominance of plants suitable for consumption (fruit trees, vegetables, and herbs), and gardens that function mainly as ornamental gardens (mostly having evergreens, ornamental plants, and tended lawns) formed separate modules. The last two modules group gardens whose owners are environmentally conscious, close to nature, and enjoy outdoor activities. In spite of the differences, inter-module connections were common, indicating that predicting gardening practices from plants grown or garden type is hardly possible. Indeed, we did not find a clear indication that the presence of certain plant types (e.g. herbs or evergreens) affects biodiversity-positive gardening. Yet, although with a low importance in the model, the presence of lawns pointed toward lower BDF scores and the presence of vegetables seemed to be related with biodiversity-positive activities. Indeed, the presence of lawns is probably related to increased herbicide use and frequent mowing whereas the presence of vegetables likely lowers pesticide input because in kitchen gardens products are grown for own consumption.

However, based on our analysis of the associations between garden characteristics and garden care practices, no associations were detected between the use of pesticides and herbicides (irrespective of the type) and any other variables. Thus, pesticide and herbicide usage appears to be random among the study population, suggesting that most domestic garden owners, regardless of socio-demographic parameters, gardening care practices, or garden parameters, use plant protection products. Indeed, although biodiversity-positive activities and garden characteristics, such as having a pond and supporting pollinators, were more typical among the respondents than those that were biodiversity-negative, more than 60% of the respondents used some pesticides (of whom almost half used synthetic pesticides) in their gardens. Albeit alarming, this is in line with the study of Varga-Szilay & Pozsgai (2022), who found excessive pesticide use even in otherwise biodiversity-friendly farmlands, as well as with a European study (Coisnon et al., 2019) in which Hungary was classified as a country least avoiding pesticide use in gardens. The latter work also pointed to the lower trust in environmental associations and the lack of reliable sources of biodiversity-related information as major culprits. Similarly to other countries, information on alternative, less harmful or biological pesticides (e.g. *Bacillus thuringiensis* or baculoviruses) is available in Hungary but the use of these is most likely linked to a narrow environmentally conscious and highly educated stratum of the society. Yet, 72% of participants in our study had higher education and access to the Internet, and thus information, which suggests that selecting and placing confidence in the available information were to be blamed. This emphasizes the significance of reaching out to garden owners through diverse channels; some may not rely on information from the Internet (Wyckhuys et al., 2019) and in these cases, knowledge transfer through personal links is likely to be the most efficient. Indeed, in our network analysis personal links and from the Internet showed a negative association. Therefore, to effectively reach such groups, prioritizing communication means relying on interpersonal interactions, such as gardening associations, community gardens, and workshops should be employed (Barthel et al., 2010). Through these channels, information can be disseminated to garden owners over 55 years old who practice conventional gardening and urban dwellers who may be more disconnected from nature. At these events, it is crucial to emphasize the harmful effects of pesticide use on individuals, the environment, and on the future of the next generations as well as to highlight non-chemical alternatives to the ones who are inclined to adopt biodiversity-friendly measures. As an added benefit, community gardens and similar shared gardening practices can also substantially improve human well-being both for isolated urban dwellers (Leavell et al., 2019) and the elderly (Fjaestad et al., 2023).

Besides pesticide use, our results highlight that mowing several times a month was also one of the common biodiversity-negative practices. For some, such as for those who own middle-size gardens, this may not be working because the limited size of these gardens potentially prevents owners from setting aside an undisturbed area. Additionally, frequent mowing of the lawn in these gardens may be more manageable, than in larger gardens (over 500m^2^). Indeed middle-sized gardens strongly and negatively affected the BDF score, most likely as a result of mowing several times a month. Explaining the time and monetary costs of frequent mowing along with the potential harm it causes to biodiversity may however convince owners to decrease mowing activities (Lerman et al., 2018). Campaigns like No Mow May can be of great help in achieving this vision. Indeed, we found that knowing about the NMMc was positively associated with active pollinator support and motivation for nature conservation. Our results also showed that knowledge of insects and observing many pollinator groups favour biodiversity-positive practices (van Heezik et al., 2012). On the other hand, we found an overlap between those who mow several times a month and those who leave unmown patches, indicating that even if gardeners can be persuaded to refrain from mowing for a short period, such as one month in the NMMc, this is probably more difficult than pursuing them to leave temporally unmown patches or undisturbed areas. Regardless, campaigns focused on biodiversity awareness hold the potential to promptly impact gardening practices, steering them towards sustainable garden care.

### 4.1. Study limits

Since in our work the respondents were not randomly selected, but rather formed a subsample of garden owners who were aware of the announcement and participated voluntarily, the study is not likely to be representative for the entire Hungarian garden owner population. Indeed, the questionnaire could only be completed online, therefore, it had a lower chance of reaching the infrastructurally less developed eastern part of the country.

In an effort to engage a broad audience through our questionnaire and to prevent potential respondents from feeling deterred by the length of the questionnaire, we attempted to make it as concise as possible, limited the number of questions, and intentionally prioritized simplicity and clarity in various instances. This sometimes meant compromising to some degree in scientific terminology and precision, for example, gardening practices, and pesticide and fertiliser use could have been recorded on a finer scale. Certain responses, inherently from the method, exhibited subjectivity. For instance, the respondents’ knowledge about their gardens’ biodiversity we relied on a self-assessed scale and the number of pollinator groups they observed in their gardens may have depended on their ecological knowledge. Nevertheless, the introduction of photo-based selection in the corresponding question probably mitigated this subjectivity.

Whilst these, and the limitations arising from the relative weaknesses of our model, may constrain the generalizability and applicability of the findings, a sufficiently large number of people completed our questionnaire, which likely ensures a thorough depiction of common gardening behaviours and trends in garden practices among household garden owners in Hungary.

## 5. Conclusion and future perspectives

The findings of our study provide a comprehensive overview on the gardening practices, motivations of garden owners, and garden characteristics in Hungary, discuss linked practices and pave the way to efficiently drive knowledge transfer and education for pursuing people toward biodiversity-friendly gardening practices. In this first step to understanding the little-studied sociological drivers of Eastern European gardening practices, we highlight the distinctions between Hungary and Western European countries and underscore the necessity for customised tactics in promoting biodiversity-friendly and sustainable gardening practices for countries with different cultural backgrounds. Due to the lack of social trust and the ageing society, this includes, for instance, favouring interpersonal programs over disseminating information online. Although our study concentrated exclusively on Hungary, it is important to recognize that these peculiarities may not be unique to this only country, but are likely to be relevant to a significant, culturally related segment of Eastern European countries. Indeed, our work could function as a novel scheme to raise studies focusing on understanding the links between garden diversities, garden characteristics and gardening practices in Eastern European countries, where environmental consciousness is expected to be lower compared to Western European countries.

However, garden biodiversity is also influenced by factors not discussed here (such as the biodiversity of the surrounding area, landscape layout, or animal behaviour (Goddard et al., 2013)). Ground truth inventories of garden biodiversities are thus also needed, along with the assessment of landscape-level variables. Citizen science projects for domestic garden (and urban greenspace) inventories provide a valuable tool both for low-cost data collection and conservation-minded education at large. Thus combining our questionnaire-based approach, with a volunteer monitoring scheme and remotely sensed datasets would be indispensable to gain vital insight for efficient conservation and maintaining biodiversity in urban areas. Yet, it is key to be mindful that a complementary interplay between social sciences and ecology is essential for substantially advancing the understanding of how the benefits of urban green spaces, including gardens, could be maximised both for conservation and human well-being.

## Supporting information

Electronic Supplementary Material

## Acknowledgements

We are grateful to Kris A. G. Wyckhuys for his valuable comments on the first version of the manuscript.

## Author information

### Author contributions

ZVS and GP conceived the idea of the project, designed the study and created the first version of the questionnaire. ZVS, GP, GS and KGF finalised the questions. ZVS and GP translated and specialised the questionnaire into English. All authors participated in data collection. Analysis was performed by ZVS and GP. The draft of the manuscript was written by ZVS and GP. All authors read and approved the final manuscript.

## Declarations

### Conflict of interest

The authors declare no competing interests.

Declaration of Generative AI and AI-assisted technologies in the writing process

#### Statement

During the preparation of this work the authors used ChatGPT (version 3.5) in order to improve readability and language. After using this tool, the authors reviewed and edited the content as needed and takes full responsibility for the content of the publication.

## Code availability

The underlying computer code is available in the GitHub repository https://github.com/zsvargaszilay/gardening_in_hungary.

