## Supplementary material for "Bridging biodiversity and gardening: Unravelling the interplay of socio-demographic factors, garden practices, and garden characteristics": Electronic Supplementary Material

Full title

Running title

Social drivers, synergies and trade-offs in biodiversity-focused gardening

Zsófia Varga-Szilay[1*], Kinga Gabriella Fetykó[2], Gergely Szövényi[3], Gábor Pozsgai[4*]

[1]Doctoral School of Biology, Institute of Biology, ELTE Eötvös Loránd University, Budapest, Hungary

[2]Independent researcher

[3]Department of Systematic Zoology and Ecology, ELTE Eötvös Loránd University, Budapest, Hungary

[4]Ce3C – Centre for Ecology, Evolution and Environmental Changes, Azorean Biodiversity Group, CHANGE – Global Change and Sustainability Institute, University of the Azores, Faculty of Agricultural Sciences and Environment, Angra do Heroísmo, Terceira, Açores, Portugal

*Corresponding authors: Zsófia Varga-Szilay, Doctoral School of Biology, Institute of Biology, ELTE Eötvös Loránd University, 1117 Budapest, Hungary,, https://orcid.org/0000-0001-9712-7654

Gábor Pozsgai, Ce3C – Centre for Ecology, Evolution and Environmental Changes, Azorean Biodiversity Group, CHANGE – Global Change and Sustainability Institute, University of the Azores, Faculty of Agricultural Sciences and Environment, Angra do Heroísmo, Terceira, Açores, Portugal,, https://orcid.org/0000-0002-2300-6558

**This file includes:**

Supplementary Figure 1-2

Supplementary Table 1-7

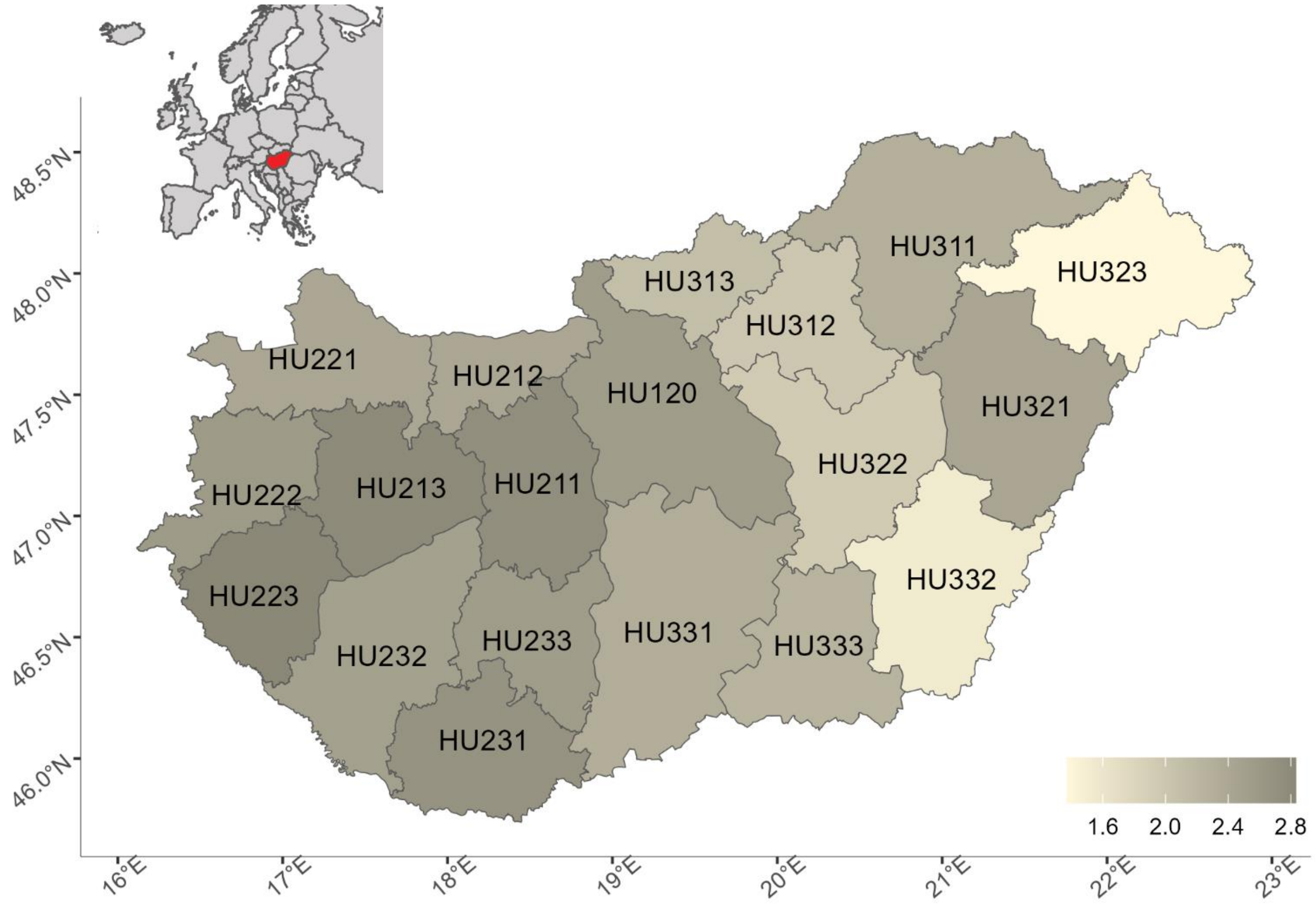


**Supplementary Figure 1**: The map shows the NUTS*-3 levels. To improve representativeness the number of respondents was standardised for 100,000 inhabitants per NUTS classification level (indicated with the colour depth in the map). The NUTS of the capital (HU110, Budapest) was merged with the largest NUTS surrounding it (HU120, Pest). The values are on a logarithmic scale. (* The Nomenclature of Territorial Units for Statistics.)

**Supplementary Table 1:** Questions, the original levels and the re-categorised levels of the questionnaire (translated from Hungarian).

| **Questions** | **Original levels** | **Re-categorised levels** |
|---|---|---|
| 1. What is the type of urban settlement where your garden is located? | Capital | City |
| | City | City |
| | Town | Town |
| | Countryside | Countryside |
| | Farmland | Countryside |
| 2. Please, provide your gender. | Male | Male |
| | Female | Female |
| | Other | Other |
| 3. Please, provide your age. | Under 18 | Excluded |
| | 18-25 | Under 36 |
| | 26-35 | Under 36 |
| | 36-45 | 36-55 |
| | 46-55 | 36-55 |
| | 56-65 | Over 55 |
| | Over 65 | Over 55 |
| 4. What is your highest level of education? | Elementary | Middle |
| | Middle | Middle |
| | Postsecondary | High |
| | Postgraduate | High |
| 5. Are there any children in your household? | Yes | Yes |
| | No | No |
| 6. What is the area of your garden? | Under 10 m2 | Small |
| | 10-50 m2 | Small |
| | 50-100 m2 | Small |
| | 100-500 m2 | Medium |
| | Over 500 m2 | Large |
| 7. Do you have a pond in your garden? | Yes | Yes |
| | No | No |
| 8. How long have you been gardening? | Less than 1 year | Newly |
| | 1 year | Newly |
| | 2-5 years ago | Newly |
| | More than 5 years, but less than 10 years | Middle |
| | More than 10 years | Long ago |

| | | |
|---|---|---|
| 9. What kind of garden do you have? Please, choose from the list what describes it the best. | House garden/Village garden<br>Flower garden<br>Kitchen garden<br>Orchard<br>Vineyard<br>Community garden<br>Other | House garden<br>Flower garden<br>Kitchen garden<br>Orchard<br>Vineyard<br>Community garden and other<br>Community garden and other |
| 10. How large area do the listed plants take up in your garden? [Vegetables; Herbs; Fruit trees; Grapes; Ornamental plants/trees; Evergreens; Lawn; Undisturbed area/Fallow] | Do not have<br>Very small<br>Medium<br>Significant<br>Most of it | No<br>Yes<br>Yes<br>Yes<br>Yes |
| 11. To what extent do the listed aspects influence your gardening habits? [Making my surroundings more beautiful; Nature conservation; Making money (e.g. with production); Self-supply/production for own use] | 1 (It does not affect at all)<br>2<br>3<br>4<br>5 (This is the most important) | No<br>Yes<br>Yes<br>Yes<br>Yes |
| 12. How do you perceive gardening? | My favourite hobby<br>A pleasant pastime<br>Opportunity to exercise<br>Duty<br>Work | Pastime<br>Pastime<br>Pastime<br>Duty<br>Duty |
| 13. Please, finish the sentence with what is the most true of you. [Plants; Birds; Insects] | I know<br>I know, and I explore them consciously<br>I do not know them well but I am trying to get to know them better<br>I do not know, but I want to know<br>I do not know | I know<br>I know<br>I do not know<br>I do not know<br>I do not know |
| 14. How often do you mow the lawn during the growing season (between April and September = 6 months)? | Several times a month<br>Once a month<br>Every two months<br>Twice<br>Once<br>I do not mow the lawn at all | Very often<br>Very often<br>Often<br>Rarely<br>Rarely<br>Never |
| 15. Do you leave unmown patches when you mow? | Yes<br>No | Yes<br>No |
| 16. How often do you plant ornamental plants? | 1 (Never) | No |

| | | |
|---|---|---|
| | 2 | Yes |
| | 3 | Yes |
| | 4 | Yes |
| | 5 (Very often) | Yes |
| 17. Where do you get/buy the seeds/seedlings/plants? | I collect the seeds from my garden | Collecting |
| | Collecting in nature in my country | Collecting |
| | Collecting in nature abroad | Collecting |
| | Shops specialised in farming and gardening | Shops |
| | Markets or producers/growers | Shops |
| | Supermarkets | Shops |
| | Webshops | Shops |
| | Seed exchange | Personal |
| | I get them from friends | Personal |
| | I get them from people who I do not know | Personal |
| 18. Do you regularly fertilize your garden? If so, what method do you use? | I use synthetic fertiliser | Synthetic |
| | I use animal manure | Non-synthetic |
| | I use compost | Non-synthetic |
| | I do not fertilize | Do not use |
| 19. Do you use herbicides containing glyphosate in your garden? | Yes, I use it | Yes |
| | I use herbicides, but I do not know the main active ingredient | Yes |
| | I use herbicides, but the main active ingredient is not glyphosate | Yes |
| | I do not use herbicides at all | No |
| 20. For what purpose do you grow plants? | Only for own use | Only for own use |
| | Both for own use and sale | Both for own use and sale |
| | Only for sale | Only for sale |
| | I do not grow plants for consumption and/or sale | I do not grow |
| 21. Do you use pesticides? If so, what type of pesticide(s) do you use? | Conventional pesticides/synthetic pesticides | Synthetic pesticides |
| | Eco/Bio/Green labelled pesticides | Eco/Bio/Green labelled pesticides |
| | Self-made pesticides, home practices | Homemade |
| | Do not use pesticides at all | Do not use pesticides at all |
| 22. What pollinators do you regularly observe/see in your garden? (The question was picture-based, with images illustrating representative members of pollinator groups commonly found in gardens.) | Butterflies | |
| | Bumblebees | "None of the above" OR less than four pollinator groups = Non or few |

| | | |
|---|---|---|
| | Honeybees | More than three AND less than six pollinator groups = Medium |
| | Other wild bees | More OR equal than six pollinator groups = Lot |
| | Wasps | |
| | Ants | |
| | Hoverflies | |
| | Other flies | |
| | Beetles | |
| | None of the above | |
| 23. How do you support wild pollinators? | Creating/preserving natural habitats (e.g. wildflower strips) | Natural habitats |
| | Creating artificial habitats (e.g. insect- or bee hotels, bumblebee homes, hoverflies-lagoons) | Artificial habitats |
| | Providing food sources (e.g. pollinator-friendly flowers) | Food sources |
| | Water sources | Water sources |
| | I do not support them actively | I do not support |
| 24. Do you think your garden is pollinator-friendly? | Yes | Yes |
| | No | No |
| 25. Can you imagine your garden being part of a garden network that helps maintain biodiversity? | Yes | Yes |
| | Maybe | Maybe |
| | No | No |
| 26. Since food sources for pollinators are the most abundant in May the #NoMowMay 2022 campaign urges people not to mow their lawns in that month. Have you heard about the campaign and if so, have you joined it? | Yes, and I did not mow in May | Yes, I joined |
| | Yes, but I did mow in May | Yes, but I did not joint |
| | I have not heard about this campaign | No |
| 27. How do you learn/gather knowledge/information about gardening? | Gardening journals/magazines, books | Traditional |
| | TV, radio | Traditional |
| | Internet | Web |
| | Social media platforms | Web |
| | Self-training groups | Personal and other |
| | Consulting with other garden owners | Personal and other |
| | Consulting with professional gardeners/agronomists | Personal and other |
| | Other | Personal and other |

**Supplementary Table 2:** Socio-demographic characteristics of the study population (n = 1260)

| | | Total (n = 1260) | |
|---|---|---|---|
| Variable | | n | % |
| Gender | Male | 343 | 27.22 |
| | Female | 916 | 72.70 |
| | Other | 1 | 0.08 |
| Age (years) | Under 36 | 246 | 19.52 |
| | 36-55 | 669 | 53.10 |
| | Over 55 | 345 | 27.38 |
| Education level | Middle | 354 | 28.1 |
| | High | 906 | 71.9 |
| Residence type | City | 392 | 31.11 |
| | Town | 342 | 27.14 |
| | Countryside | 526 | 41.75 |
| Children | Yes | 616 | 48.89 |
| | No | 644 | 51.11 |

**Supplementary Table 3**: Results of ANOVA-like permutational test for the db-RDA investigating the relationships between garden characteristics, gardening practices and socio-demographic variables.

| Variables | Sum Of Sqs | Pr(>F) |
|---|---|---|
| Type of residence | 1.384 | 0.001* |
| Gender | 0.828 | 0.001* |
| Age | 2.581 | 0.001* |
| Education level | 0.565 | 0.001* |
| Having children | 0.210 | 0.003* |
| Residual | 114.080 | |

**Supplementary Table 4**: Presence (%) of the biodiversity-positive (A) and biodiversity-negative (B) gardening practices among the study population (n = 1260).

| **A** | Yes | | No | |
|---|---|---|---|---|
| Variable | n | % | n | % |
| Water sources | 870 | 69.05 | 390 | 30.95 |
| Flower sources | 828 | 65.71 | 432 | 34.29 |
| Natural habitats | 827 | 65.63 | 433 | 34.37 |
| Leave unmown patches | 786 | 62.38 | 474 | 37.62 |
| Do not use pesticides | 477 | 37.86 | 783 | 62.14 |
| Artificial habitats | 423 | 33.60 | 837 | 66.40 |
| Jointed to NoMowMay | 284 | 22.54 | 976 | 77.46 |
| Have pond | 184 | 14.60 | 1076 | 85.40 |

| **B** | Yes | | No | |
|---|---|---|---|---|
| Variable | n | % | n | % |
| Lack of undisturbed area | 636 | 50.48 | 624 | 49.52 |
| Mowing several times a month | 404 | 32.06 | 856 | 67.94 |
| Use synthetic pesticides | 340 | 26.98 | 920 | 73.02 |

| | | | | |
|---|---|---|---|---|
| Use herbicides | 190 | 15.08 | 1070 | 84.92 |
| Use synthetic fertilisers | 158 | 12.54 | 1102 | 87.46 |

**Supplementary Table 5:** Presence (%) of herbicide usage among the study population (n = 1260).

| Age | Do you use herbicides containing glyphosate in your garden? | n | Percentage of age per usage category (%) | Percentage of usage category per age (%) |
|---|---|---|---|---|
| Under 36 | Yes, I use it | 23 | 9.35 | 17.6 |
| | I use herbicides, but the main active ingredient is not glyphosate | 8 | 3.25 | 30.8 |
| | I use herbicides, but I do not know the main ingredient | 8 | 3.25 | 24.2 |
| | I do not herbicides at all | 207 | 84.2 | 19.4 |
| 36-55 | Yes, I use it | 73 | 10.9 | 55.7 |
| | I use herbicides, but the main active ingredient is not glyphosate | 9 | 1.35 | 34.6 |
| | I use herbicides, but I do not know the main ingredient | 18 | 2.69 | 54.6 |
| | I do not herbicides at all | 569 | 85.0 | 53.2 |
| Over 55 | Yes, I use it | 35 | 10.1 | 26.7 |
| | I use herbicides, but the main active ingredient is not glyphosate | 9 | 2.61 | 34.6 |
| | I use herbicides, but I do not know the main ingredient | 7 | 2.03 | 21.2 |
| | I do not herbicides at all | 294 | 85.2 | 27.5 |

**Supplementary Table 6**: Relative influence of factors generated from the Gradient Boosting Machine (GBM) model (R-squared: 0.16, RMSE: 0.32) for predicting Biodiversity-friendliness score (BDF). Variables marked with asterisk (*) have two levels.

| Variables | Relative influence |
|---|---|
| Observing many pollinators groups* | 11.6765120 |
| Garden type | 9.1441346 |
| Motivation: nature conservation* | 8.4094770 |
| Information through personal link* | 7.4622973 |
| Knowledge about insects* | 5.5137041 |
| Garden size | 5.3844236 |
| Age | 4.7598829 |
| Duration of gardening experience | 4.7029874 |
| Having herbs* | 4.4805255 |
| Type of residence | 4.2282597 |
| Getting seeds/plants via collection* | 4.1335063 |
| Having ornamental plants* | 3.9911554 |
| Knowledge about plants* | 3.1628923 |
| Getting seeds/plants in person* | 2.7812964 |
| Growing crops* | 2.6756280 |
| Planting ornamental plants* | 2.5842374 |
| Garden perception* | 2.3670376 |
| Having children* | 2.0329137 |

| | |
|---|---|
| Education level* | 1.6659437 |
| Information from the Internet* | 1.4621913 |
| Having grapes* | 1.4114943 |
| Knowledge about birds* | 1.2887135 |
| Having vegetables* | 1.1260483 |
| Having evergreens* | 0.8103696 |
| Having lawn* | 0.7386529 |
| Gender | 0.6975678 |
| Information through traditional media* | 0.4750882 |
| Observing several pollinator groups* | 0.3088561 |
| Having fruit trees* | 0.2923316 |
| Motivation: production* | 0.2318714 |
| Motivation: beautiful garden* | 0.0000000 |

**Supplementary Table 7:** Results of the ANOVA-like permutational test for the RDA investigating the relationships between garden characteristics, gardening practices and socio-demographic variables.

| Variables | Variance | Pr(>F) |
|---|---|---|
| Type of residence | 1.0884 | 0.060**.** |
| Gender | 0.3572 | 0.776 |
| Age | 0.9193 | 0.445 |
| Education level | 0.6776 | 0.035* |
| Having children | 0.4458 | 0.546 |
| Residual | 5.0322 | |

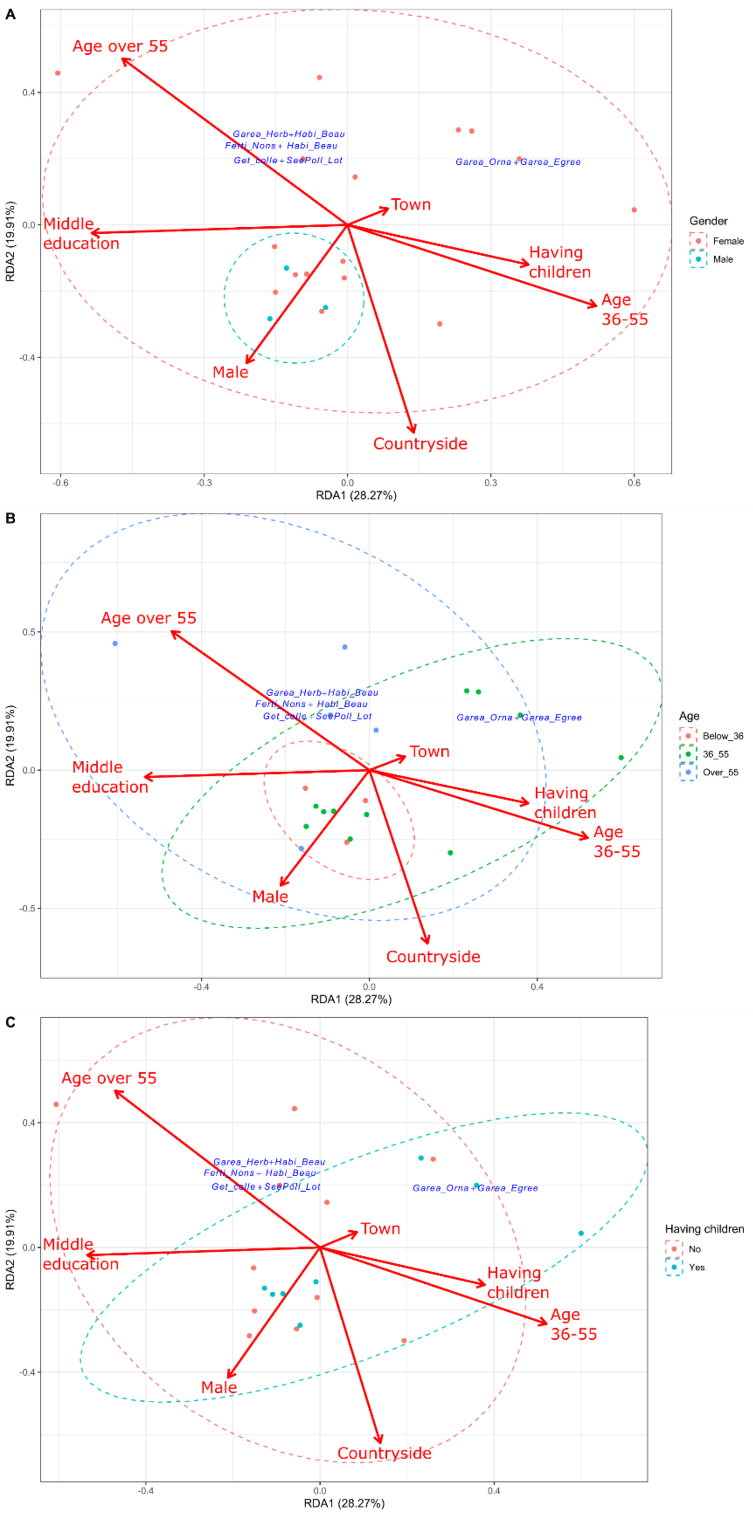


**Supplementary Figure 2:** Redundancy analysis (RDA) plot showing the associated pairs of gardening practices and garden characteristics, along with the explanatory socio-demographic variables. The length and direction of the vectors represent the strength and direction of the relationship. The ellipses represent the 95% confidence intervals of associations of the gender (A), the age (B), and having children (C). Only associations (blue) whose RDA scores on the first two axes were lower than the mean of zero and the smallest value on the axes or greater than the mean of zero and the greatest value on the axes are shown. (Abbreviations: Garea_Herb: having herbs, Habi_Beau: motivation: beautiful garden, Ferti_Nons: use non-synthetic fertilisers, Get_colle: getting seeds/plants via collection, SeePoll_Lot: observing many pollinators groups, Garea_Orna: having ornamental plants, Garea_Egreev: having evergreens.)
